# Dynamic microtubules slow down during their shrinkage phase

**DOI:** 10.1101/2022.07.27.501773

**Authors:** Anna Luchniak, Yin-Wei Kuo, Catherine McGuinness, Sabyasachi Sutradhar, Ron Orbach, Mohammed Mahamdeh, Jonathon Howard

## Abstract

Microtubules are dynamic polymers that undergo stochastic transitions between growing and shrinking phases. The structural and chemical properties of these phases remain poorly understood. The transition from growth to shrinkage, termed catastrophe, is not a first-order reaction but is rather a multi-step process whose frequency increases with the growth time: the microtubule ages as the older microtubule tip becomes more unstable. Aging shows that the growing phase is not a single state but comprises several substates of increasing instability. To investigate whether the shrinking phase is also multi-state, we characterized the kinetics of microtubule shrinkage following catastrophe using an *in vitro* reconstitution assay with purified tubulins. We found that the shrinkage speed is highly variable across microtubules and that the shrinkage speed of individual microtubules slows down over time by as much as several fold. The shrinkage slowdown was observed in both fluorescently labeled and unlabeled microtubules as well as in microtubules polymerized from tubulin purified from different species, suggesting that the shrinkage slowdown is a general property of microtubules. These results indicate that microtubule shrinkage, like catastrophe, is time-dependent and that the shrinking microtubule tip passes through a succession of states of increasing stability. We hypothesize that the shrinkage slowdown is due to destabilizing events that took place during growth which led to multi-step catastrophe. This suggests that the aging associated with growth is also manifest during shrinkage with the older, more unstable growing tip being associated with a faster depolymerizing shrinking tip.

**Statement of Significance:** The dynamics of the microtubule cytoskeleton is crucial for several functions in eukaryotic cells. Microtubule dynamics is traditionally described by constant growth and shrinkage speeds with first order transitions between the growth and shrinkage phases. However, catastrophe, the transition from growth to shrinkage, increases with microtubule age and is not a first order process. In contrast to the common assumption that microtubules shrink with constant speed, here we show that shrinking microtubule tips undergo step-wise slowdown during depolymerization. Our results suggest that microtubule shrinkage, like catastrophe, is a multi-step process. This finding is important for understanding the molecular nature of microtubule dynamic instability and how microtubule shrinkage can be modulated by microtubule associated proteins.

## Introduction

Microtubules, essential components of the eukaryotic cytoskeleton, are cylindrical filaments made up of typically 13 protofilaments. They are highly dynamic polymers that stochastically switch between phases of slow growth and fast shrinkage (1). The switch from growth to shrinkage is termed catastrophe and is thought to be due to the loss of a cap of GTP-tubulin at the growing end (2). The switch from shrinkage to growth is termed rescue and is thought to be due to reincorporation of GTP-tubulin during the shrinking process (3, 4). These transitions, termed dynamic instability, control turnover of microtubule polymers (5, 6) and allow microtubule tips to explore cytoplasmic space to capture chromosomes during mitosis (7), to create pushing forces that position the mitotic spindle (8) and the nucleus (9), and to fill axons and dendrites during neuronal morphogenesis (10–12).

The simplest model of dynamic instability posits that the growing and shrinking phases are each single states and that the transitions between them are first order (13). This model predicts the existence of a critical concentration above which growth is unbounded and below which growth is bounded, and the mean length is determined by the growth and shrinkage speeds and the transition rates (14, 15). Despite these successful predictions, the two-state model has been challenged by several experimental observations. For example, the growth speed is not constant: during the growth phase, the length does not increase steadily but fluctuates on a sub-second timescale with an amplitude much larger than that expected for the random association and dissociation of single tubulin subunits to the growing end (16–19). Furthermore, the growth speed decreases prior to catastrophe (20, 21). Thus, the growth phase is not a single state, but includes several substates.

Catastrophe is also a complex dynamical process. Once deemed to be a single-step stochastic event with a constant frequency over time, the collection of large datasets showed that catastrophe is instead a multi-step process that depends on the age of the microtubule, with older microtubules having higher catastrophe frequencies (22, 23). It was proposed that the growing tip incorporates three or more defects (analogous to the accrual of multiple mutations in cancer development) that increase the likelihood of the tip undergoing a catastrophe (analogous to the onset of a cancer) (22, 24). A potential structural basis of multi-step catastrophe is the evolution of the tip structure, a form of memory (25, 26). For example, randomly occurring lattice defects, such as a change in protofilament number (27) or lattice-start number (28) may irreversibly alter the structural state of the growing tip making it more prone to catastrophe. Thus, length-dependent catastrophe, in addition to the growth speed fluctuations, also indicates that the growing tip changes its properties (i.e., state) over time.

Unlike growth and catastrophe, the shrinkage phase after catastrophe is poorly characterized. During shrinkage, protofilaments peel and curl outwards, and tubulin dissociates from the shrinking protofilament tips, causing the rapid shortening of the filament (29). The average microtubule shrinking speeds reported for microtubules assembled *in vitro* from mammalian brain tubulin vary widely, typically between 10 and 40 μm/s (30–32). Because microtubule shrinking events are short-lived, the accuracy of these measurements is limited by the temporal resolution of imaging. In our study, we have circumvented this limitation by using interference-reflection microscopy (IRM) to image microtubule shrinkage with high temporal resolution. We show that the shrinkage speed is highly variable from microtubule to microtubule and does not depend on the polymer’s length at catastrophe. Furthermore, as individual microtubules shorten, the depolymerization speed slows down up to several-fold compared to the initial shrinkage speed. We hypothesize that microtubule shrinkage, like catastrophe, is a multi-step process, and that changes in shrinkage speeds may correspond to alterations in lattice stability that occurred during growth.

## Results

To examine the kinetics of microtubule shrinkage following catastrophe, we visualized the dynamic microtubules *in vitro* using interference-reflection microscopy (IRM) (Figure 1A) (33). Stabilized microtubule seeds polymerized from TAMRA-labeled tubulin in the presence of the slowly hydrolysable GTP analog GMPCPP were attached to the surface via anti-TAMRA antibodies. Dynamic microtubules were then grown from these stabilized seeds by introducing unlabeled bovine-brain tubulin in the presence of GTP. To capture the fast depolymerization process, we imaged the growth and shrinkage of dynamic microtubules with high frame rate (8-10 Hz). Surprisingly, while some microtubules shrank with constant speed (Figure 1B, left), a large fraction initially shrank quickly but then slowed down as they approached the seed (Figure 1B, 1C). The variation in the shrinkage profiles was not seed-dependent, as we observed separate shrinkage events from the same microtubule seeds, some having constant speed and others slowing down (Figure 1B red and yellow boxes). By tracking the position of depolymerizing tips during the course of shrinkage (Figure S1), we found that the majority of the shrinkage events showed faster initial shrinkage speeds than their final speeds (Figure 1D), with a decrease of 4.1 ± 2.7 fold (mean ± SD, *N* = 91 microtubules from 3 experiments). This suggests that microtubules frequently decelerate during depolymerization. The mean shrinking speed of individual microtubules was highly variable (Figure 1E; 0.388 ± 0.144 μm/s (mean ± SD, *N* = 91 microtubules), consistent with the shrinkage speed slowing down in a stochastic manner.

**Figure 1.**
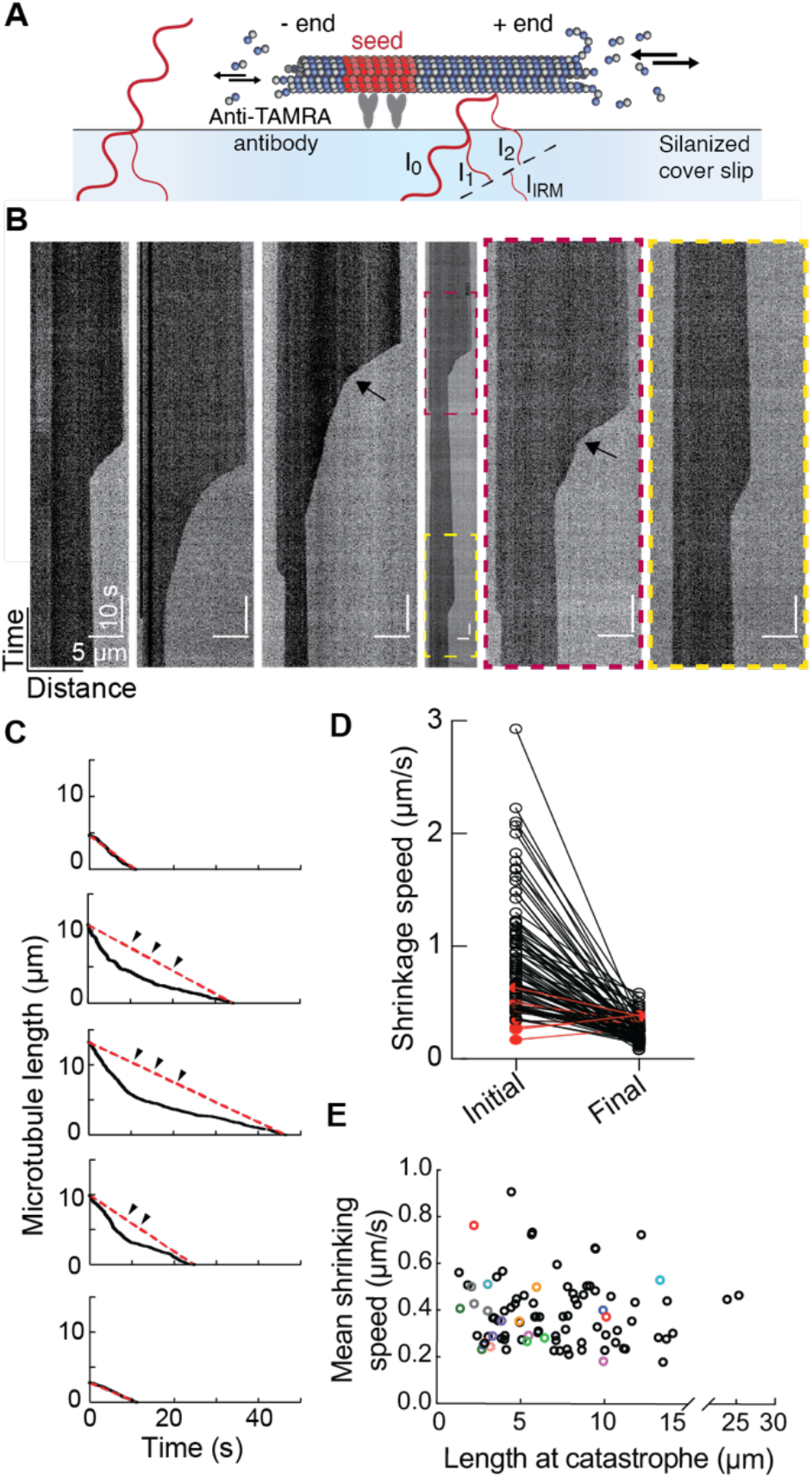
The shrinking speed of *in vitro* microtubules is highly variable. (A) Schematic of *in vitro* dynamic microtubule assay. Label-free microtubules were grown from TAMRA-labeled GMPCPP-stabilized microtubule seeds attached to a silanized glass surface using anti-TAMRA antibodies. Dynamic microtubules were imaged by time-lapse interference-reflection microscopy (IRM) in which light reflected from the glass/water interface interferes with that reflected at the water/microtubule interface. I_IRM_: interference intensity; I_0_: intensity of incident light; I_1_: intensity of light reflected of glass/sample interface; I_2_: intensity of light reflected from water/microtubule interface. (B) Representative kymographs depicting shrinkage profiles of individual microtubule polymers (microtubules corresponding to the dark regions). Maroon and yellow dashed boxes mark shrinkage events of microtubules growing from the same seed. Horizontal scale bar: 5 μm, vertical scale bar: 10 seconds. Example of clear deceleration events are indicated by arrows. (C) Microtubule depolymerization curves corresponding to kymographs in panel (B). Red dashed lines represent hypothetical depolymerization tracks for microtubules disassembling with a constant speed, matching the mean shrinking speed of a microtubule depicted in the graph. Black arrowheads highlight regions where the depolymerization speed deviates from the mean shrinking speed of a given microtubule. (D) Comparison of initial and final shrinking speed for bovine microtubules. Initial speed was calculated as a mean shrinking speed over the initial 10% of microtubule length starting from the position of microtubule tip at the time of catastrophe. The final speed is the mean shrinking speed over the final 10% of the dynamic microtubule extension prior to reaching the microtubule seed. Microtubule length is defined as the length of the dynamic microtubule extension at catastrophe. Points labeled in red represent microtubules without shrinkage slowdown whose average decelerations were not significantly different from zero (8 out of 91 microtubules; see text description of Fig. 3C for details). (E) Mean shrinking speed of microtubules is independent of microtubule length at catastrophe (*N*=91 events from triplicate experiments). Shrinkage events of dynamic microtubules originated from the same seeds were highlighted with the same colors.

To bolster our confidence that the slowdown is a general property of microtubules, we did two control experiments. First, fluorescently labeled microtubules imaged by total-internal-reflection fluorescence (TIRF) microscopy also showed deceleration during shrinkage (Figure 2A, S3), showing that the observed shrinkage slowdown was also present for labeled tubulin and not an artifact of the IRM imaging. Second, to test whether the deceleration of shrinkage requires the presence of free GTP-tubulin, we severed GMPCPP-tubulin capped GDP-microtubules with a severing enzyme, spastin (34), in the absence of free tubulin, and imaged shrinking microtubules (Figure S2A). The severed microtubules also experienced slowdown of depolymerization (Figure S2B-S2D), suggesting that the deceleration of shrinkage is a feature inherent to the microtubule lattice and does not require the addition of free tubulin onto the shrinking microtubule tips.

**Figure 2.**
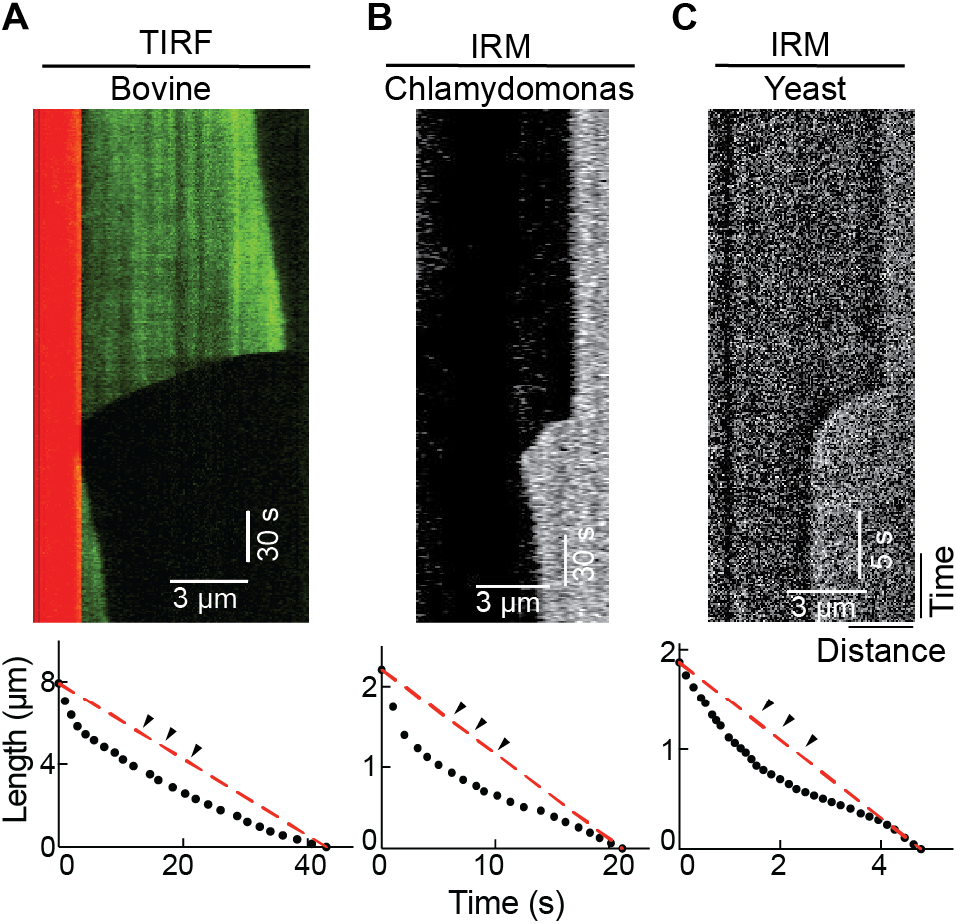
Slowdown is observed for bovine, *Chlamydomonas* and yeast microtubules. (A) Representative kymograph of microtubule shrinkage visualized using TIRF microscopy with the corresponding depolymerization curve. (B,C) Kymographs and matching plots of microtubule shrinkage for microtubules assembled from unlabeled tubulin isolated from *Chlamydomonas reinhardtii* and budding yeast and imaged using IRM. Red dashed line indicates position of the microtubule tip for a filament disassembling at the rate matching mean shrinking speed of the microtubule in the graph. Black arrowheads highlight regions where microtubule shrinking speeds deviate from mean depolymerization speed.

We next asked whether microtubules polymerized from tubulin purified from different organisms exhibit similar shrinkage slowdown. We imaged dynamic microtubules polymerized from budding-yeast tubulin (35) and *Chlamydomonas* axonemal tubulin (36) using IRM. In both cases, we observed shrinkage events with fast initial depolymerization that slowed over time (Figure 2B, 2C), similar to the depolymerization curves of microtubules polymerized from bovine brain tubulin (Figure 1C). Thus, dynamic microtubules assembled from tubulin purified from various sources displayed slowdown of shrinkage after catastrophe.

We next quantified the change of shrinkage speed during depolymerization. The instantaneous shrinkage speed of bovine microtubules changed rapidly over time (Figure 3A). The large change in the shrinkage speed is reflected by the high standard deviation of the instantaneous shrinkage speed 0.313 ± 0.163 μm/s, which is larger than our estimated detection accuracy (0.063 μm/s, see Methods) and the variability in instantaneous shrinkage speed showed no evident correlation with the length at catastrophe (Figure 3B). To quantify shrinkage slowdown, the instantaneous speed was fitted with a linear regression model where the slope indicates the average shrinkage deceleration. The bovine brain microtubules decelerate 0.046 ± 0.073 μm/s^2^ (mean ± SD, Figure 3C, imaged by IRM), which is significantly larger than zero (two-tailed one sample t-test, N=91 events, p<0.0001). Similar to the bovine brain tubulin (Figure 3C), deceleration of the shrinkage rate was observed in microtubules assembled from yeast (Figure 3D, S4A and C) and *Chlamydomonas* axonemal tubulin (Figure S4B, D). The magnitude of deceleration showed no evident correlation with the filament length at catastrophe (Fig. 3E, S4A). Interestingly, yeast microtubules showed a larger slowdown (Figure 3D), potentially due to the structural differences between yeast and mammalian microtubules (37). These results demonstrate that most dynamic microtubules experience shrinkage slowdown during depolymerization.

**Figure 3.**
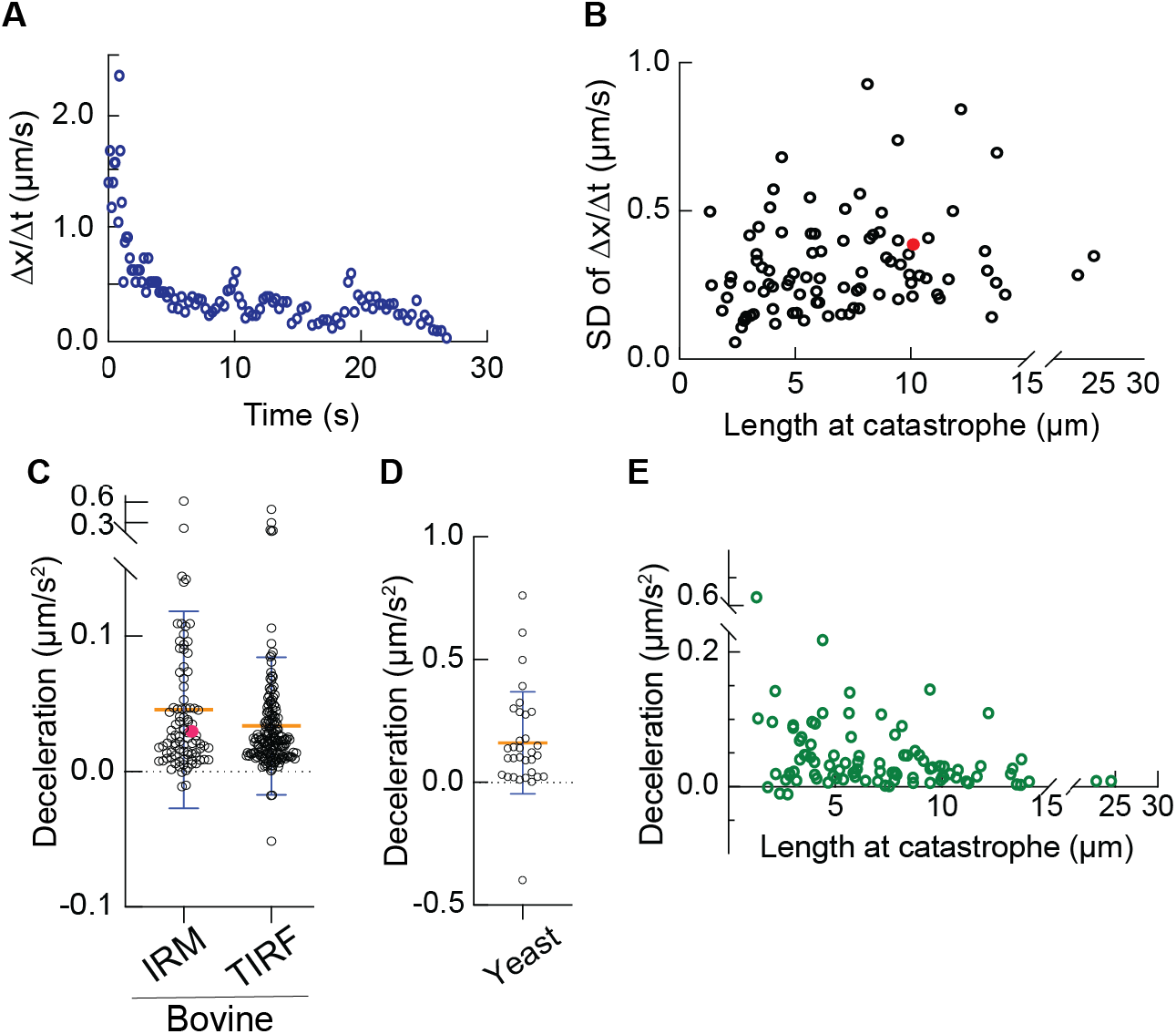
Microtubules decelerate during shrinkage. (A) Instantaneous speed of microtubule shrinkage (*dx/dt*) is variable during microtubule shortening. (B) Standard deviation of instantaneous speed of microtubule shrinkage is independent of polymer’s length at catastrophe (N = 91 microtubules). The point labeled in red corresponds to the standard deviation of instantaneous speed of shrinkage for the microtubule presented in panel A. (C,D) Deceleration of shrinking speed *d^2^x/dt^2^* for microtubules assembled from bovine (N_IRM_ = 91, N_TIRF_ = 163) and yeast tubulin (N = 32). The point labeled in magenta corresponds to deceleration of the microtubule presented in panel A (error bars: mean ± SD). (E) Deceleration of bovine microtubules (imaged by IRM) showed no evident correlation with the length at catastrophe. Coefficient of determination of linear regression R^2^ = 0.07 and slope= - 0.0046 (s^−2^).

Many microtubule depolymerization traces showed clear transitions from higher to lower speeds (see example in Figure 1B, arrows), suggesting that depolymerization of microtubules may slow down in a stepwise manner instead of by a continuous, gradual deceleration. We fitted the depolymerization traces with a piecewise-linear function, varying the number of segments (an example of such a fit is shown in Figure 4A). As expected, the fit improved as the number of segments increased (Figure 4B). The root-mean-square error decreased to below 0.1 μm when three or more segments were used. Thus, similar to catastrophe, the microtubule shrinkage is also consistent with a multistep process, with three steps capturing 99.1% of the variance (Figure S5).

**Figure 4.**
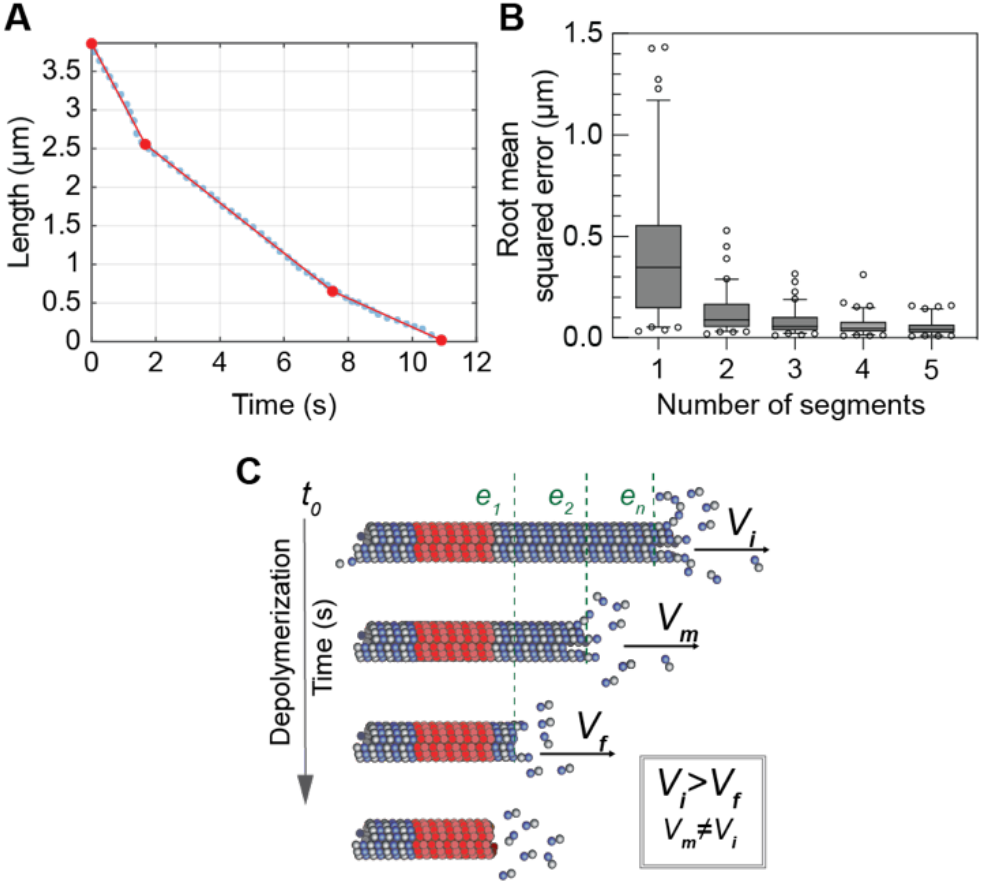
Microtubule shrinkage *in vitro* is a multistep process. (A) A microtubule depolymerization trace (dotted line) fitted with 3 segments using piecewise-linear function (red lines between the red points). (B) Root-mean-square error of the piecewise linear fit decreased below 0.1 μm when 3 or more segments were used. The horizontal lines indicate the 75^th^, 50^th^, and 25^th^ percentiles; whiskers: 5^th^, 95^th^ percentile. N=91 shrinkage events. (C) Model of microtubule shrinkage as a multi-step process. Growing microtubule encounters stochastic destabilizing events (e_1_, e_2_, e_n_), that are embedded as a ‘memory’ into microtubule lattice (green dashed lines). Accumulation of three events results in a microtubule catastrophe with a transition from growth to shrinkage. The highly unstable microtubule lattice, which has experienced multiple destabilizing events, has the highest shrinking speed (*V_i_*). When a depolymerizing microtubule tip passes one region, in which the ‘memory’ of a destabilizing encounter is stored, to the next, the microtubule lattice becomes more stable and the shrinking speed decreases (*V*_m_, *V*_f_). Consequently, the most stable microtubule lattice is found in the proximity of the microtubule seed, and in that region, microtubule shrinking speed is slow (*V*_f_). Thus, stochastic events experienced by a microtubule during its growth alter the stability of the microtubule lattice. Accordingly, microtubule shrinkage can be considered a multi-step process and divided into steps characterized by different shrinking speeds and stability of the polymer’s lattice.

## Discussion

Using an *in vitro* reconstitution system, we discovered that microtubules do not depolymerize at a constant speed, as implicitly assumed in the literature when a shrinkage rate is reported. Instead, the shrinkage speed of microtubules frequently slows down over time. We do not believe that the shrinkage slowdown is caused by contamination of microtubule-associated proteins (MAPs) as no visible contaminants were observed in the bovine, yeast (Figure S6) or *Chlamydomonas* (36) tubulin preparations. Moreover, shrinkage slowdown was observed in microtubules polymerized from bovine, yeast and *Chlamydomonas* tubulin; because each tubulin was purified using a different method, it is unlikely that they all contain a common contaminant. Finally, we were able to identify multiple examples of deceleration during microtubule depolymerization in the published work from various labs (38–44), either with tubulin alone or with different types of MAPs. Thus, we conclude that the slowdown of shrinkage is likely to be an intrinsic property of dynamic microtubules.

What is the mechanistic origin of the shrinkage slowdown after catastrophe? The depolymerization traces are consistent with three or more shrinkage phases with decreasing velocities (Figure 4B), suggesting that the shrinking tip starts from a highly unstable state after catastrophe (which depolymerizes quickly) and becomes more stable over time (depolymerizes more slowly). This process and its multistep nature are reminiscent of the aging of microtubules during their growth (22, 25, 26, 45), where three steps are required to lead to catastrophe (Figure 4B). We therefore propose a model where several stochastic destabilization events accumulate during the polymerization phase, eventually leading to a catastrophe. The lattice contains the “memory” of these destabilization events, and we propose that the depolymerizing tip “reads” these memories in a reverse order during depolymerization. The shrinking microtubule tip thus encounters regions of the lattice with increasing stability, which effectively slows down microtubule shrinkage (Figure 4C). This model implies that the destabilization “steps” that induce catastrophe alter the microtubule lattice, perhaps in the form of structural inhomogeneity. Recent optical and electron microscopy methods that facilitate the detection of subtle structural changes of the tips or along the microtubule wall (28, 46–48) may provide novel insights into the molecular basis of these destabilization steps and how the shrinking microtubule tip structure evolves during slowdown.

An interesting avenue to explore is the role of the carboxy-terminal tail (CTT) in shrinkage slowdown. Removal of the negatively-charged CTT from β-tubulin results in a slower average shrinking speed, which is likely due to the reduction of the electrostatic repulsion and therefore increase in the affinity of tubulin to the microtubule tip (49). Removal of β-tubulin’s CTT also causes a higher catastrophe frequency (49). This may be due to the accumulation of lattice irregularities, as tubulin without CTTs has a higher tendency to assemble into aberrant structures such as sheets, spirals and twisted filaments (50, 51). Whether microtubules assembled from tubulin without CTTs also exhibit a larger shrinkage slowdown remains to be determined.

Another interesting topic to explore is the relationship between shrinkage slowdown and rescue. Several rescue-promoting MAPs have been found to decrease the average shrinkage speed: yeast CLASP (52) (but not human CLASP (53, 54)), tau (55), spastin (56) and TPX2 (57). Lattice islands containing GMPCPP that promote rescue also showed a slower depolymerization rate (58). It is thus tempting to speculate that rescue factors may suppress the initial fast shrinkage phases during depolymerization. The change in depolymerization rate also implies that, similar to catastrophe, rescue may not be a single-step random process either and the probability of rescue may increase over time due to the slower shrinkage speed. Notably, recent studies have shown that repeated rescues can occur at distinct regions of the lattice, frequently associated with damage and structural irregularities of the lattice and their repair (24, 27, 31, 59, 60). Examining if the shrinkage rate also slows down at these rescue hotspots may indicate whether rescues promoted by these lattice irregularities or by rescue-promoting proteins share similar features. Our knowledge about the molecular nature of catastrophe and rescue is still very limited. Scrutiny of the microtubule depolymerization process can potentially offer new information regarding the biophysical principles underlying microtubule dynamic instability.

## Materials and Methods

### Protein expression and purification

Bovine tubulin was purified using two cycles of polymerization and depolymerization in high molarity buffer from bovine brains as previously described (61). Purified tubulin was aliquoted, flash frozen in liquid nitrogen and stored at −80°C until use. Tubulin from *Chlamydomonas reinhardtii* was purified as described in (62).

Buddying yeast tubulin was purified as described before using TOG column (35, 39). Briefly, *S. cerevisiae* (strain BY4741) was grown at 30°C to a logarithmic growth phase. Cells were pelleted, washed with doubly distilled water and ground in liquid nitrogen as previously described (35). Cell powder was resuspended in the lysis buffer (80 mM K-PIPES, 1 mM EGTA, 1 mM MgCl_2_, pH 6.9, 10% glycerol, 0.2% Triton X-100, 1 mM DTT, 0.1 mM ATP, 1 mM GTP, 0.2 mM Pefabloc, 5 μg/mL leupeptin, 1 μg/mL pepstatin A, 1 μg/mL aprotinin, Benzonase 125 units/ml). The lysate was stirred for 10 min at 4°C and cleared via centrifugation at 125000 g for 40 min at 4°C. The supernatant was collected, filtered through 0.45μm filter, de-gassed for 15 minutes and applied onto TOG-column pre-equilibrated with the lysis buffer. The column was washed with wash buffer (80mM K-PIPES, 1mM EGTA, pH 6.9, 100μM GTP, 0.1%Tween, 5mM ATP, 10 mM MgCl_2_). Tubulin was eluted with linear gradient over 20 column volumes (Buffer A: 80mM K-PIPES, 1mM EGTA, 1mM MgCl_2_, pH 6.9; Buffer B: 750 mM (NH_4_)_2_SO_4_, 80mM K-PIPES, 1mM EGTA, 1mM MgCl_2_, pH 6.9). Purified yeast tubulin aliquoted and snap frozen in liquid nitrogen in 80 mM K-PIPES, 1 mM EGTA, 1 mM MgCl_2_, pH 6.9, 10% glycerol.

Short (208 aa to end) isoform of *Drosophila melanogaster* spastin was expressed in bacteria (Rosetta-DE3-competent cells; Novagen), purified and stored as previously described (56).

### Dynamic microtubule assay

The dynamic microtubule assays were carried out in microchannels prepared from two silanized coverslips sealed with parafilm (63). GMPCPP-stabilized microtubule seeds labeled with TAMRA dye (64) were used to initiate the growth of dynamic microtubule extensions. To measure shrinkage of bovine microtubules, 7.5 μM bovine unlabeled GTP-tubulin was used while 4 or 5 μM yeast tubulin was used to polymerize dynamic yeast microtubules. The dynamics of unlabeled microtubules was visualized using interference reflection microscopy (33, 65) in the assay buffer (80 mM K-PIPES, 1 mM EGTA, 1 mM MgCl_2_, pH 6.9, 1 mM DTT, 2 mM GTP). All time-lapse images were collected with a sCMOS camera (Zyla 4.2, Andor, Belfast, Scotland) mounted on a Ti Eclipse Nikon microscope using a 100x/1.49 NA Apochromat objective (Nikon, Melville, NY). The IRM dynamic assays were imaged at 10 Hz rate for 10 minutes at 34°C for bovine, or 28°C for yeast and Chlamydomonas microtubules.

The *in vitro* dynamic microtubule assays with labeled tubulin were conducted in microchannels as described above. Microtubules were assembled from bovine, Alexa Fluor 488-labeled, GTP-tubulin at 8 μM concentration in assay buffer, supplemented with oxygen scavenger mix (40 mM glucose, 40 μg/mL glucose oxidase, 16 μg/mL catalase, 0.1 mg/mL casein, 10 mM DTT). Dynamics of labeled microtubules was visualized using total internal reflection microscopy (63) at 1 Hz frame rate for 10 minutes at 34 °C.

### GMPCPP-capped microtubules severing assay

GMPCPP-stabilized microtubule seeds polymerized from biotinylated and TAMRA-labeled bovine tubulin were attached to the surface of a microchannel using anti-biotin antibodies (Sigma-Aldrich). Microtubules were assembled using 12 μM unlabeled bovine GTP-tubulin for 10 min at 28 °C in polymerization buffer (80 mM K-PIPES, 1 mM EGTA, 1 mM MgCl_2_, pH 6.9, 5 mM DTT, 1 mM GTP). Subsequently, 5 μM TAMRA-labeled tubulin in polymerization buffer containing 0.5 mM GMPCPP was flowed into the reaction channel to cap and stabilize microtubule filaments for 5 minutes. The positions of microtubule seeds and GMPCPP-tubulin caps were recorded using TIRF microscopy prior to the addition of spastin. 1 nM spastin in severing buffer (80 mM K-PIPES, 1 mM EGTA, 1 mM MgCl_2_, pH 6.9, 1 mM DTT, 1 mM ATP, 50 mM KCl) was added into the reaction channel, and microtubule shrinkage following severing at 28 °C was visualized using IRM (65) and recorded at 10 Hz.

### Image analysis

Fiji software was used to analyze microtubule shrinking velocities. Briefly, kymographs of individual microtubules were generated following the subtraction of background IRM images, generated by averaging images of empty reaction channels from the corresponding time series to enhance contrast. The average microtubule shrinkage rates were calculated based on the ratio of horizontal and vertical distances on the kymograph, corresponding to the distance of a microtubule tip at catastrophe from the seed and time spent in shrinking phase, respectively. To track position of microtubule tip over time during shrinking events, kymographs generated from IRM time-lapse images were filtered using Gaussian blur to reduce random noise and thresholded. Subsequently, the boundary of a microtubule on the kymograph was found using edge detection function and applied as a mask on top of a corresponding, unprocessed kymograph to facilitate microtubule tip tracking (Figure S1). Kymographs generated from TIRF time-lapse images were subjected to the same process prior to tracking of microtubule tip during shrinkage, except Gaussian blur step was omitted. The detection accuracy was estimated by the standard deviation of the tracked length of GMPCPP-stabilized microtubules without dynamic microtubule extension over several seconds.

### Quantification of mean and instantaneous shrinkage speed

The mean shrinkage speed and uncertainty in the speed for all analyzed samples was calculated from linear regression fits of microtubule position as a function of time during shrinkage events. The microtubule tracking data was decimated to verify there is no correlation between adjacent points and that the number of independent data points within the set was not overestimated. The mean speed and variance were compared with the mean speed and variance of decimated data sets (Table S1).

The instantaneous speed consists of the points Δ*x_n_* = (*x*_*n*+1_ – *x_n_*)/Δ*t* measured at times, *t_n_* = *n* · Δ*t* where Δ*t* is the time interval of the video and *n* = 0, 1, 2, … *N* – 1 is the sample number. The instantaneous speed data was fitted with linear regression to calculate the average deceleration and uncertainty in deceleration.

### Piece-wise linear curve fitting

A MATLAB based piecewise linear fitting algorithm known as the ‘SLM-Shape Language Modeling’ (66), which has also been employed recently to understand dendritic tip dynamics (67), is used to fit microtubule depolymerization curves. We used a variable number of segments with the positions of the segment ends as free variables to fit the data. The root mean squared error decreases with increasing number of segments as shown in Figure 4B.

### Statistical analysis

Statistical analysis for all individual experiments and repeats was performed using GraphPad Prism7 or MATLAB statistical software. Evaluation of statistical significance is described in the text and respective figure legends.

## Author Contributions

A.L. and J.H. conceived the project, A.L., Y.-W.K. performed the bovine tubulin experiments, C. M. performed the yeast tubulin experiments, R.O. performed the *Chlamydomonas* tubulin experiments, M.M. assisted the development of high-speed imaging and image analysis pipeline, S.S. performed the piece-wise linear fit, A.L., Y.- W.K. and J.H. wrote the manuscripts with input from all authors.

## Acknowledgements

The authors thanks Dr. Maijia Liao, Dr. Olivier Trottier and Dr. Pei-Zhi Ivy Huang for the fruitful discussions during the development of the project. R.O. was supported by cross-Disciplinary Fellowship from Human Frontier Science Program (LT000919/2015-C) and European Molecular Biology Organization long-term fellowship (ALTF 1424-2014). This work was supported by NIH Grant R01 GM139337 and R01 NS118884 (to J.H.)

