## Supplementary information for "Dynamic microtubules slow down during their shrinkage phase"

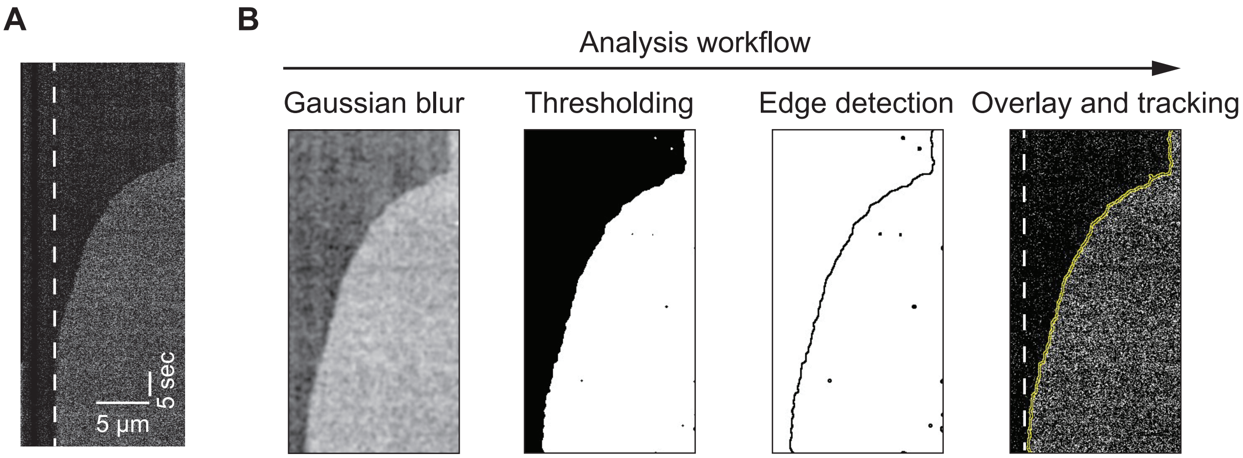


**Figure S1. Analysis workflow of microtubule shrinkage**

(A) Kymograph depicting shrinkage of bovine microtubule visualized by IRM. Dashed line indicates position of microtubule seed. (B) Preparation of kymographs for analysis of microtubule shrinkage. A Gaussian filter was applied to reduce image noise. Next, kymograph was thresholded to generate binary image for microtubule detection. Identified microtubule outline was applied on top of the original kymograph to facilitate detection of microtubule position over time. Subsequently position of microtubule tip during shrinkage was tracked and recorded to generate microtubule depolymerization curves.

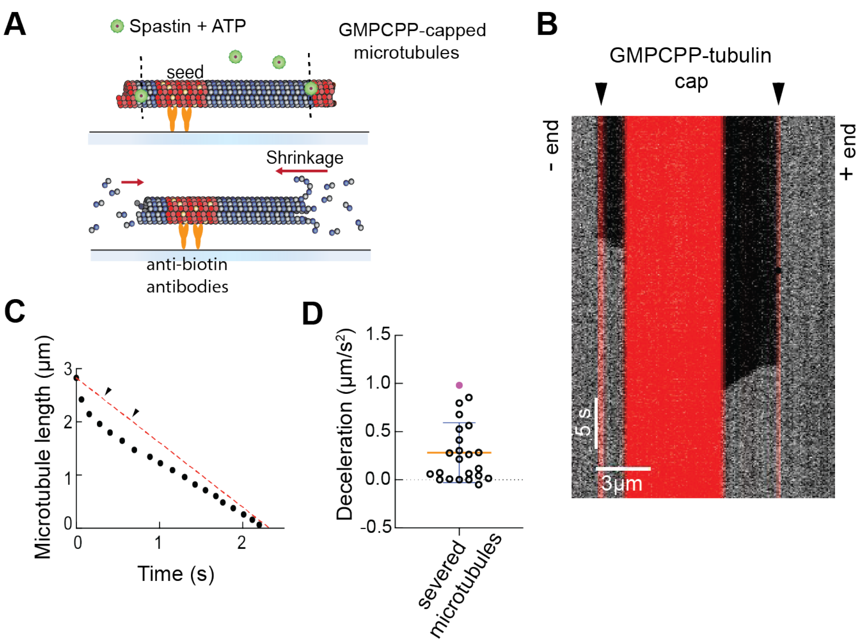


**Figure S2. Shrinkage slowdown of bovine microtubules in the absence of free tubulin**

(A) Schematic of microtubule depolymerization assay in the absence of free tubulin. Label-free microtubules were grown from biotinylated, TAMRA-labeled GMPCPP-stabilized microtubule seeds attached to a silanized glass surface using anti-biotin antibodies. To create GMPCPP-stabilized cap at the ends of dynamic microtubules, unlabeled free tubulin was washed away from the reaction channel and replaced with a buffer containing TAMRA-labeled tubulin and GMPCPP. The addition of fluorescently-labeled tubulin to the ends of dynamic microtubules was verified by TIRF microscopy. Finally, excess of GMPCPP and labeled tubulin was removed and 1 nM spastin with 1 mM MgATP was used to sever microtubule polymers (indicated as black dashed lines). Severing and depolymerization of microtubules was visualized by time-lapse IRM. Only microtubules severed close to the fluorescently-labeled cap were analyzed in this assay. (B) Representative kymograph of microtubule shrinkage in the absence of free tubulin. Black arrowheads indicate the position of GMPCPP-stabilized, TAMRA-labeled microtubule cap. (C) Microtubule depolymerization curve corresponding to kymograph in panel B. Red dashed line indicates position of microtubule tip for a polymer disassembling at the rate matching the mean shrinking speed of the microtubule in the graph. (D) Shrinking speed of bovine microtubules decelerates during depolymerization in the absence of free tubulin (N = 23; Mean ± SD). Point highlighted in magenta reflects deceleration of microtubule presented in the kymograph in panel B.

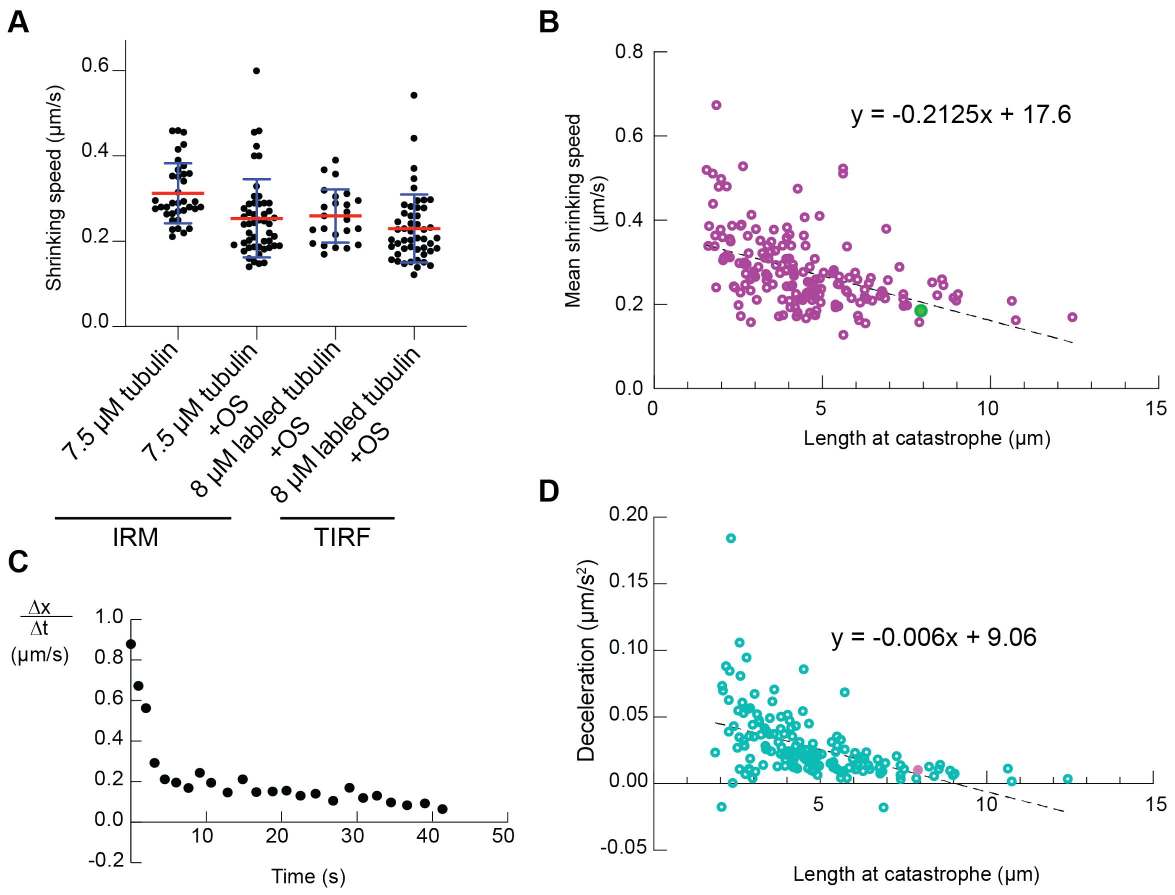


**Figure S3. Bovine microtubules imaged by TIRF microscopy decelerate during shrinkage**

(A) Mean shrinking speed for microtubules visualized using different imaging techniques. Shrinking speed was measured from the kymograph. Specifically, it was calculated from the slope of a line drawn along the depolymerizing microtubule. 7.5 µM tubulin (N = 36 events, n = 3 experiments); 7.5 µM tubulin + OS (N = 53, n = 3); 8 µM labeled tubulin + OS (N = 24, n = 2); 8 µM labeled tubulin + OS imaged by TIRF microscopy (N = 49 n = 4). OS: oxygen scavenger mix (Mean ± SD). (B) Mean shrinking speed of bovine microtubules visualized by TIRF microscopy shows weak correlation with the microtubule length at catastrophe ($p<0.0001$, F test, N = 176), potentially due to the slow accumulation of photodamage. Black dashed line represents linear regression with the coefficient of determination ($R^2$) 0.22. Point colored in green corresponds to mean shrinking speed of the microtubule from Fig 2A. (C) Shrinking speed is highly variable during microtubule shortening. Plot depicting change in instantaneous speed during microtubule shrinkage corresponds to the microtubule from Fig 2A (D) Shrinking speed of bovine microtubules decelerates during microtubule shortening. Black dashed line represents linear regression with the coefficient of determination ($R^2$) 0.24 (N = 163). Point colored in magenta depicts deceleration of the microtubule from Fig 2A.

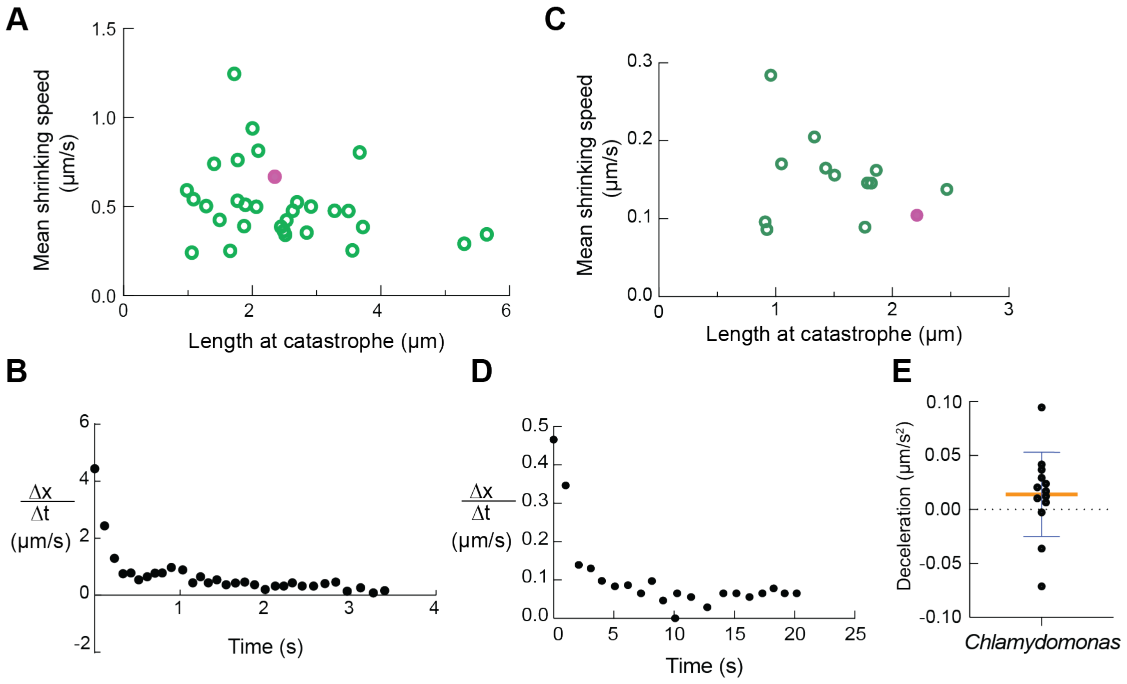


**Figure S4. Deceleration of shrinking speed for yeast and *Chlamydomonas* microtubules**

(A) Mean shrinking speed of yeast microtubules does not depend on the microtubule length at catastrophe. Point colored in magenta corresponds to mean shrinking speed of the microtubule presented in Fig 2C. Black dashed line represents linear regression with the coefficient of determination ($R^2$) = 0.08 and slope = -0.0559 (1/s). (B) Mean shrinking speed for microtubules assembled from *Chlamydomonas* tubulin is independent of microtubule length at catastrophe. Point colored in magenta corresponds to mean shrinking speed of the microtubule presented in Fig 2B. Black dashed line represents linear regression with the coefficient of determination ($R^2$) 0.06 and slope -0.027 (1/s). (C)(D) Instantaneous speed of microtubule shrinkage (dx/dt) is variable during disassembly of yeast (C) and *Chlamydomonas* (D) microtubules. Instantaneous speed plots correspond to microtubules depicted in Fig 2C and B, respectively. (E) Average shrinkage rate deceleration of microtubules assembled from *Chlamydomonas* tubulin.

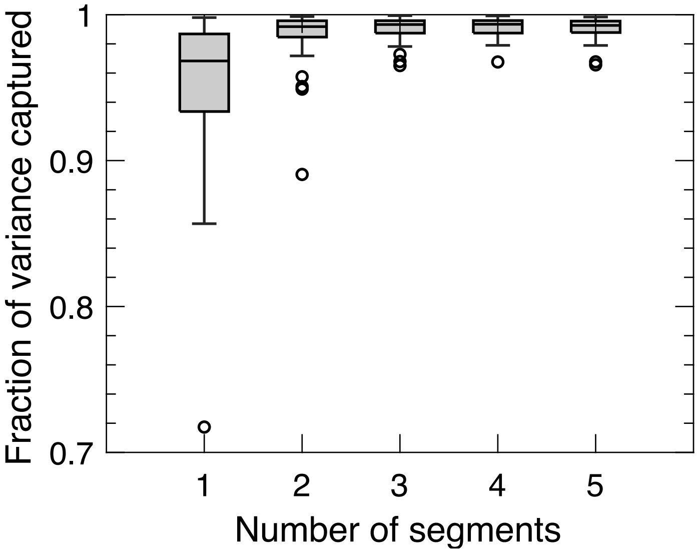


**Figure S5. The fraction of variance captured ($R^2$ value) by the piecewise linear fitting as a function of number of segments used to fit the microtubule depolymerization curves**

The fraction of variance captured increases with number of segments (95.5% for 1 segment, 98.8 % for 2 segments and 99.1% for 3 or more segments) and saturates after 3 or more segments (also see Figure 4A, B in the main text).

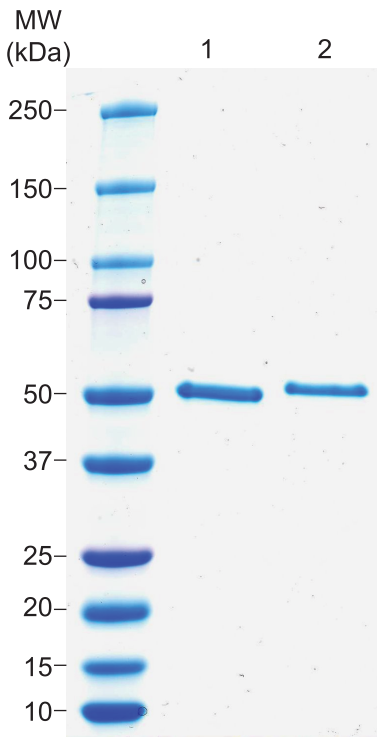


**Figure S6. Purification of bovine and yeast tubulin**

Coomassie-stained SDS-PAGE gel loaded with molecular weight standards (MW) tubulin purified from bovine brain (1) and *Saccharomyces cerevisiae* (2).

| MT | $\sigma_v^2$ n=0,1,2,3,…*N* | $\sigma_{v2}^2$ n=0,2,4,…*N* |
|---|---|---|
| 1 | 9.23667E-06 | 1.98459E-05 |
| 2 | 0.000261455 | 0.000587494 |
| 3 | 9.27847E-06 | 2.00236E-05 |
| 4 | 3.69426E-05 | 7.6308E-05 |
| 5 | 4.54459E-06 | 9.27092E-06 |
| 6 | 0.000128154 | 0.000291596 |
| 7 | 4.81543E-06 | 9.92507E-06 |
| 8 | 1.73046E-05 | 3.5923E-05 |
| 9 | 1.39704E-06 | 2.87256E-06 |
| 10 | 6.85563E-05 | 0.000147935 |
| 11 | 1.17738E-05 | 2.44444E-05 |
| 12 | 7.07775E-06 | 1.43986E-05 |
| 13 | 3.8553E-05 | 9.83983E-05 |
| 14 | 0.002226801 | 0.005096742 |
| 15 | 5.85094E-06 | 1.18395E-05 |
| 16 | 3.74339E-05 | 7.66587E-05 |
| 17 | 0.000232797 | 0.000492299 |
| 18 | 2.54028E-05 | 5.25366E-05 |
| 19 | 1.41045E-05 | 2.95996E-05 |
| 20 | 0.000174855 | 0.000357733 |
| 21 | 0.000109386 | 0.000226972 |
| 22 | 2.17929E-05 | 4.49147E-05 |
| 23 | 2.56709E-05 | 5.28483E-05 |
| 24 | 8.25134E-06 | 1.63313E-05 |
| 25 | 1.48617E-05 | 3.10592E-05 |
| 26 | 5.13176E-05 | 0.000103895 |
| 27 | 5.22417E-05 | 0.000112286 |
| 28 | 5.20234E-05 | 0.000122483 |
| 29 | 9.36957E-05 | 0.000191499 |
| 30 | 0.000103352 | 0.000206352 |
| 31 | 2.64717E-05 | 5.50288E-05 |
| 32 | 7.82801E-06 | 1.63748E-05 |
| 33 | 4.56385E-05 | 9.53408E-05 |
| 34 | 4.51102E-05 | 9.31074E-05 |
| 35 | 1.20934E-05 | 2.52861E-05 |
| 36 | 7.35331E-05 | 0.000163598 |
| 37 | 2.33657E-06 | 4.89127E-06 |
| 38 | 7.80461E-06 | 1.58343E-05 |
| 39 | 5.70802E-06 | 1.20358E-05 |
| 40 | 2.1487E-05 | 4.39816E-05 |
| 41 | 3.99906E-05 | 8.15792E-05 |
| 42 | 9.0758E-06 | 1.87706E-05 |
| 43 | 3.25564E-06 | 6.935E-06 |
| 44 | 7.85849E-06 | 1.62445E-05 |
| 45 | 4.90713E-06 | 1.01194E-05 |
| 46 | 3.47571E-06 | 7.09761E-06 |
| 47 | 8.65414E-05 | 0.000180754 |
| 48 | 6.0872E-05 | 0.000128771 |
| 49 | 0.000272593 | 0.000573745 |
| 50 | 0.000103576 | 0.000220621 |
| 51 | 6.80181E-05 | 0.000141909 |
| 52 | 0.00030006 | 0.000643481 |
| 53 | 9.97617E-05 | 0.000210804 |
| 54 | 0.00013521 | 0.000284944 |
| 55 | 5.80708E-05 | 0.000123108 |
| 56 | 0.000407442 | 0.000892978 |
| 57 | 6.64318E-06 | 1.52999E-05 |
| 58 | 1.3959E-05 | 2.91676E-05 |
| 59 | 6.95364E-05 | 0.000106168 |
| 60 | 7.83419E-05 | 0.000171348 |
| 61 | 3.17832E-05 | 7.88812E-05 |
| 62 | 0.000120815 | 0.000310593 |
| 63 | 0.000289066 | 0.000599217 |
| 64 | 5.25022E-05 | 0.000101851 |
| 65 | 0.000703313 | 0.001473847 |
| 66 | 4.23781E-05 | 8.81312E-05 |
| 67 | 1.2706E-05 | 2.63595E-05 |
| 68 | 8.33532E-05 | 0.000182472 |
| 69 | 3.88558E-06 | 8.50815E-06 |
| 70 | 1.32775E-05 | 2.66592E-05 |
| 71 | 0.000897036 | 0.0018725 |
| 72 | 3.68522E-05 | 7.8779E-05 |
| 73 | 0.000350999 | 0.000592188 |
| 74 | 0.000310369 | 0.000670045 |
| 75 | 1.8315E-05 | 3.69328E-05 |
| 76 | 0.000146357 | 0.00030643 |
| 77 | 0.000281994 | 0.00061632 |
| 78 | 6.75234E-05 | 0.000143783 |
| 79 | 0.0001732 | 0.000350985 |
| 80 | 0.000335345 | 0.000738072 |
| 81 | 0.001291205 | 0.002805023 |
| 82 | 0.000764597 | 0.001580655 |
| 83 | 6.37758E-05 | 0.000131077 |
| 84 | 5.07981E-06 | 1.19618E-05 |
| 85 | 9.45556E-06 | 2.036E-05 |
| 86 | 3.56353E-05 | 7.40526E-05 |
| 87 | 0.000136347 | 0.000281852 |
| 88 | 1.31891E-05 | 2.56972E-05 |
| 89 | 1.63078E-05 | 3.57113E-05 |
| 90 | 3.96745E-05 | 9.56602E-05 |
| 91 | 2.13566E-05 | 4.17988E-05 |

**Table S1. Estimation of variance of mean shrinking speed for complete and decimated data sets.** To test for correlation between adjacent points, microtubule depolymerization data was fitted with linear regression to estimate mean shrinking speed and its variance ($\sigma_v^2$) for each analyzed microtubule. Data was subsequently decimated and only every other point from the data set was taken to calculate shrinkage speed and its new variance ($\sigma_{v2}^2$). Variance of shrinking speed of decimated data is approximately twice the variance of full data set for analyzed microtubules ($\sigma_{v2}^2 \approx 2\sigma_v^2$). Consequently, there is no correlation between the adjust points in the data sets, and microtubule shrinkage was not oversampled.
